# Disparities in avian genomic resources highlight gaps across the Global South

**DOI:** 10.64898/2026.11.05.764686

**Authors:** K. L. Vinay, Chiti Arvind, Naman Goyal, Ashwin Warudkar, Sushma Reddy, V. V. Robin

## Abstract

Avian genomes have proliferated rapidly in recent years while expanding our knowledge of genomic evolution. However, evaluating bird sampling across the tree of life and across their distributions is necessary to assess their contributions to global bird diversity and diversification patterns. We evaluated all available avian genome assemblies across geography, taxonomy, conservation status, and time. The 1,712 species with at least one genome assembly represent only ∼15% of recognized avian diversity. Although all orders were represented, 24 bird families (all restricted to the Global South) lacked any genome assembly, and only ∼37% (N= 2,397) of recognized genera had genomic representation. Nearly 78% of the most evolutionarily distinct and globally endangered (EDGE) lineages lacked a genome. Genomic resources are concentrated in temperate regions of the Global North, whereas species-rich tropical regions in Africa, South America, and Southeast Asia remain severely underrepresented relative to their diversity. Even after accounting for conservation status, evolutionary distinctiveness, and ecology, Global South species had roughly fourfold lower odds of having a genome than Global North species. Notably, the post-2021 surge in genome sequencing has counter-intuitively widened this gap. Together, our results highlight persistent gaps in genomic resources across the Global South and the need to collectively address these global knowledge gaps via collaborative efforts.

## Introduction

Global diversity follows a strong gradient, with the tropics harboring the overwhelming majority of Earth’s biodiversity (Myers et al., 2000) while the temperate, nearctic and palearctic are less species rich. This uneven distribution is exemplified by the world’s biodiversity hotspots, concentrated within tropical latitudes (Figure 1a, adapted from Hoffman et al., (2016)). These biodiversity rich regions often lack conservation resources thus creating a void. This undersampling has been previously reported for of tropical clades in comparative studies (Reddy 2014), as well as for reference genomes of tetrapods (Linck & Cadena, 2024), reptiles, and amphibians (Carneiro et al., 2025). In this context, the taxa that contribute most to global diversity contribute least to the datasets from which broad evolutionary and conservation inferences are drawn. These same regions also face the most acute pressure from land-use change and climate change (Borges et al., 2019; Jetz et al., 2007; Stork et al., 2009) resulting in ‘coldspots’ in genomic resources where conservation needs are highest (Carneiro et al., 2025).

**Figure 1:**
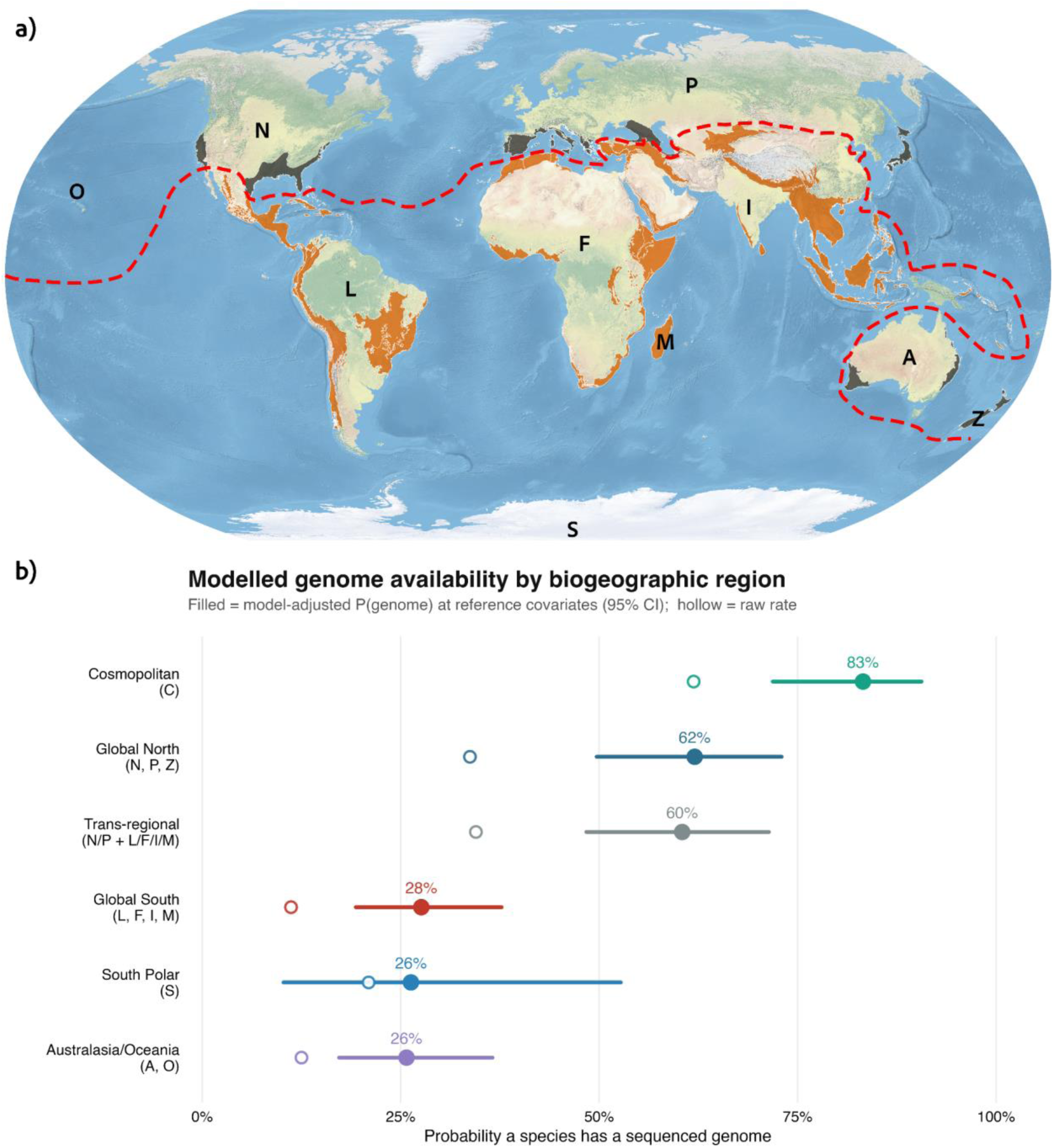
Global Distributions of the biodiversity hotspots and genome resource availability probability. a) Map showing the 36 currently recognized biodiversity hotspots as defined by (Hoffman et al., 2016). Only hotspot areas are displayed, with outer limits omitted for visual clarity. Hotspots are classified by their occurrence in the Global North (grey) and the Global South (orange) divided by the dotted line. The Mediterranean Basin hotspot spans both regions, and its extent is correspondingly divided to reflect this classification. Major biogeographic realms defined in BirdBase are denoted by letters: A - Australian; F-Afrotropical; I - Indomalayan; L - Neotropical; M - Madagascar & islands; N - Nearctic; O - Oceania; P - Palearctic; S - South Polar; Z - New Zealand & islands. b) Model-adjusted probability of genome availability by biogeographic region. Filled points show model-adjusted probability of genome availability (mixed-effects logistic regression; reference covariates; 95% CI); hollow points show raw, unadjusted rates. Regions are ordered by adjusted probability.

The geography of these biodiversity hotspots maps closely onto the geopolitical landscape. The Global South (GS) and the Global North (GN) are categorised based on the geopolitical and socio-economic groupings of states (Heath, 1981; Odeh, 2010), with the Global South including the Neotropical, Afrotropical, Indomalayan, and parts of the Australasian biogeographic realms. Although long used in geopolitics and development policy, the concept of the Global South has gained considerable traction in recent years in academia (Bull & Banik, 2025; Haug et al., 2021). Beyond its geopolitical context, the Global South is characterized by shared histories of colonization, economic vulnerability, and challenges in scientific capacity-building and policy implementation (Aregbeshola & Adekunle, 2024; Barros Leal Farias, 2024; Mormina & Istratii, 2021).

Historic and contemporary global economic conditions result in multiple inequalities. These include the lack of domestic funding for basic research, uneven knowledge production (Chankseliani, 2023), and a dependency on external, high-resource countries (often in the Global North) for advanced technologies and expertise. In the genomics era, the same broad pattern is exacerbated (Carneiro et al., 2025; Linck & Cadena, 2024). High-throughput sequencing centers and computing facilities are concentrated in Global North. The establishment and maintenance of such cutting-edge infrastructure is resource intensive and makes it hard for researchers in the Global South to break into this field. The modern genomic disparity may also have stemmed from long-standing historical patterns of access and collection of specimens (Ball et al., 2025), and research funding (Ball et al., 2025; Petersen, 2021; Rakotonarivo & Andriamihaja, 2023). Field explorations over 200 years have led to the collection of millions of specimens primarily stored and studied in natural history museums in Europe and North America (Johnson et al., 2023; Mills & Lassiter, 2026; Park et al., 2023). This results in many projects continuing to be led by institutions in the Global North, even when the focal species are endemic to the Global South. This can reinforce a form of ‘parachute’ or ‘colonial’ science (Asase et al., 2022), where samples are collected in biodiverse-rich regions and sequenced abroad with minimal local involvement in study design, analysis, or authorship (de Vos, 2020; de Vos & Schwartz, 2022).

The huge increase in genomic resources mirrors a broader shift across biodiversity research, where high-throughput sequencing is increasingly seen as a standard tool and has even fueled conservation science (Cassin-Sackett et al., 2019; Lerner & Fleischer, 2010; Oyler-McCance et al., 2016). Genome assemblies and comparative datasets are now used not only to resolve deep evolutionary relationships but also to understand fine-scale population genetic structure, map quantitative traits, and inform management plans for threatened species (Forester & Lama, 2022; Hendricks et al., 2018; Sagar et al., 2021). As these resources become foundational rather than optional, differences in the capacity to generate them are increasingly reflected in which taxa and regions are represented in our understanding of biodiversity, and in which ones can benefit from genomically informed conservation.

### Study System and Current Status

Birds, with over 11,000 recognized species, are the most diverse group of tetrapods and have played a key role in evolutionary biology research (Brusatte et al., 2015; Rheindt et al., 2025), from early ideas on natural selection to contemporary work on understanding phylogenomics and speciation (Feng et al., 2020; Jarvis et al., 2014; Joseph, 2018). Over the last decade and a half, multiple coordinated projects like Bird 10K Genomes (B10K) (Zhang et al., 2015), the Earth BioGenome Project (EBP) (Lewin et al., 2018), Genomes 10K, (Genome 10K Community of Scientists, 2009), Vertebrate Genomes Project (VGP) (https://vertebrategenomesproject.org/), Darwin Tree of Life Project (Darwin Tree of Life Project Consortium, 2022) and related consortia have produced thousands of genomes facilitating research on phylogenetics, genomic evolution, functional mechanisms, and demographic history. By 2021, these efforts had yielded roughly 500 avian genomes, a tremendous advancement that included approximately 5% of extant bird diversity (Bravo et al., 2021). These high-quality assemblies require high-molecular-weight DNA from fresh or cryopreserved tissue and blood and are likely skewed towards species in the Global North, where established tissue collections and low barriers to fieldwork make such sampling easier. Since then, genomic technologies have become more accessible and less expensive, making the generation of new genomes tenable at the population level. However, how genome production over the past five years has been distributed across regions and taxa remains unclear. Here, we examine the current distribution of avian genomic resources across the globe, with a particular focus on representation of the Global South. We specifically ask: a) How are the patterns of genome sequencing of avian species distributed across the globe? Does genome richness mirror species richness, and is the Global South underrepresented? b) Which orders and families are best represented in terms of genome assemblies, and is there a taxonomic skew across the avian tree of life? and c) How have genome representation patterns changed over time?

## Methods

### Taxonomy

We followed the Handbook of the Birds of the World-BirdLife International’s v10 (BLI hereafter) taxonomy (HBW and BirdLife International, 2025), as these have species distribution range maps (BirdLife International and Handbook of the Birds of the World Version 2024). We manually matched species names to the BLI nomenclature before proceeding with our analysis.

### Data Compilation

To quantify the global distribution in genomic resources for birds, we assembled a consolidated species-level dataset by integrating i) a genome assembly list for birds from the National Center for Biotechnology Information (NCBI), ii) BirdBase v2025 - an avian biogeographic and trait dataset, and iii) the HBW/BirdLife International species range map with BLI v10 checklist.

We downloaded a list of all the available genome assemblies for birds using ‘aves’ as a keyword on the NCBI genome page (https://www.ncbi.nlm.nih.gov/datasets/genome/) as of 06 March 2026, which returned 2,902 genome assemblies. We cleaned the list downloaded from the NCBI records to remove duplicate records (GenBank and RefSeq copies of the same assembly), leaving 2,712 assemblies, and removed one interspecific hybrid. When multiple genome versions (and alternate haplotypes) existed for a species, we retained only the most recent assembly, based on NCBI’s reported release date, to facilitate the capture of long-read information where available, leaving 1,781 genomes. Because only a small number of assemblies were designated with subspecies information, all assemblies were reduced to retain only species names, thereby avoiding counting of multiple assemblies of the same species; this removed another 36 assemblies. After this filtering, species names were matched against the BLI v10 checklist, and all mismatches were manually reviewed: of the 103 mismatched names, 70 could be reliably assigned to a BirdLife species, and assemblies that could not be matched were discarded, yielding a final dataset of 1,712 species. Because the sequencing technology reported in the NCBI assembly page is inconsistent, and no standardized classification exists, we manually categorized the assembly as either short-read, long-read, or unknown based on the metadata given on the assembly page.

Trait datasets are critical aspects for advancing biological research and can inform global conservation targets. We utilized BirdBase (Şekercioğlu et al., 2025) to obtain a wide range of ecological, life-history, and biogeographical traits for all bird species. To assess the genomic resource coverage for threatened and evolutionarily distinct species, we also obtained the latest EDGE2 scores (McClure et al., 2023) and IUCN categories.

### Species Range and Spatio-temporal Analysis

To assess the geographic distributions, we used expert-drawn range maps from BLI. We restricted the polygons to breeding and resident (season codes 1 and 2) ranges to reflect each species’ core geographic extent. Because meaningful comparisons across the globe benefit from an equal-area framework (Feeman, 2000), all range data were projected into an equal-area coordinate system. We then generated 100 km equal-area cells on a shapefile of Earth’s landmasses from rnaturalearth v1.1.0 (Massicotte & South, 2025). Only terrestrial grid cells were retained for further analysis. This grid served as a framework for estimating species richness and the distribution of taxa with genomes. We then calculated the richness as the number of distinct species overlapping in each cell.

To evaluate the temporal patterns in genomic representation, we further divided the list of species with genomes into two groups: 1) genome assemblies released before January 2021, and 2) genome assemblies released afterward. Within each grid cell, we calculated the number of species with genomes and the proportion of local richness represented by these species separately for each group, so that the difference between the two periods shows where coverage has grown and where it has not. We then compared genomic coverage between the Global North and the Global South. Because species richness differs substantially between the two regions, raw counts would reflect that difference rather than coverage. For each grid cell we therefore calculated the proportion of that cell’s resident species with a genome assembly, and summarized these per-cell proportions as a richness-weighted mean per region (GN or GS), weighting each cell by the total number of species in that cell. This was done separately for the pre- and post-2021 groups. Final richness and proportional coverage were reprojected to Robinson coordinates for visualization. All spatial processing was conducted using the *sf* package (Pebesma, 2018) in R version 4.4.1 (R Core Team, 2024).

### Modeling Genome Resource Availability

To test whether genomic resource availability differs by biogeographic region while accounting for other trait-based correlates of sequencing effort, we modeled genome availability as a binary using a generalized linear mixed model (GLMM) with a binomial error distribution and logit link. The biogeographic region for all the species was derived from the BirdBase (Şekercioğlu et al., 2025) realm classification. Because many species occupy more than one realm, each species was classified using a full set of realm assignments rather than a single dominant realm category. Species coded as Cosmopolitan formed their own category, as did all remaining species occurring in the South Polar realm. Species occurring in both a Global South realm (Neotropical, Afrotropical, Indomalayan, Madagascan or Wallacean) and a Global North realm (Nearctic or Palearctic), along with species coded as Eastern Hemisphere, were classified as Trans-regional. Of the remaining species, those occurring in the Global South realm were classified as Global South, and those occurring in the Nearctic or Palearctic, together with species restricted to New Zealand, were classified as Global North. Species restricted to the Australasian and/or Oceanian realms, including those also occurring in New Zealand, formed a separate Australasia/Oceania category, because these realms include countries on both sides of the Global North–South divide.

We included covariates expected to influence sequencing probability independently of region. Conservation status was represented as a three-level factor (Non-threatened [LC/NT], Threatened [VU/EN/CR], and Extinct [EX/EW]) based on IUCN Red List categories; Data Deficient (DD) species were excluded because they lack a defined threat classification. We additionally included EDGE species, reflecting the targeted sequencing of evolutionarily distinct, globally endangered lineages by conservation-genomics initiatives. We retained the “Island endemism” classification as is. Primary diet and primary habitat were taken from BirdBase’s weighted classifications and collapsed into broader classes to avoid sparsely populated categories. Diet was grouped into seven guilds: Invertivore (BirdBase categories Invertebrate, Beeswax), Frugivore (Fruit), Granivore (Seed), Nectarivore (Nectar), Herbivore (Herbivore, Plant), Omnivore (Omnivore), and Carnivore (Vertebrate, Fish, Carnivore, Scavenger, Ovivore). Habitat was grouped into five classes: Forest (Forest), Woodland (Woodland), Open (Grassland, Savanna, Shrub, Desert, Plains, Rocky), Aquatic (Wetland, Riparian, Coastal, Sea), and Other (Artificial, Bamboo). Invertivore and Forest were used as reference levels, being the most species-rich category in each case.

To account for phylogenetic non-independence and clade-level variation in sequencing effort, we included nested random intercepts from taxonomic order and family within order. Models were fitted by maximum likelihood with the Laplace approximation using the lme4 package (Bates et al., 2015) in R version 4.4.1 (R Core Team, 2024), with the BOBYQA optimizer (Powell, 2009). The overall contribution of the biogeographic region was tested with a likelihood-ratio test comparing the full model to a reduced model omitting the region term. Variance explained was summarized using marginal and conditional R²GLMM (Nakagawa et al., 2017; Nakagawa & Schielzeth, 2013), and model discrimination was assessed using the area under the receiver operating characteristic curve (pROC package; Robin et al., 2011). To confirm that the estimated regional effect was not an artifact of the taxonomic random-effect specification, we refit the model with a single random intercept for order only and confirmed the Global South effect estimate was materially unchanged.

## Results

Our final dataset included 1,712 species with at least one genome assembly, representing approximately 15.3% of all recognized bird species. Only 878 out of 2,397 recognized genera are currently represented with genomes, leaving close to 63.4% of the genera without any genomes. Among species with genome assemblies, long-read-based assemblies accounted for 18.17% (311), short-read assemblies for 81.60% (1,397), and 0.23% (4) were classified as unknown due to incomplete or missing metadata reporting of the sequencing technology (Figure 2A).

**Figure 2:**
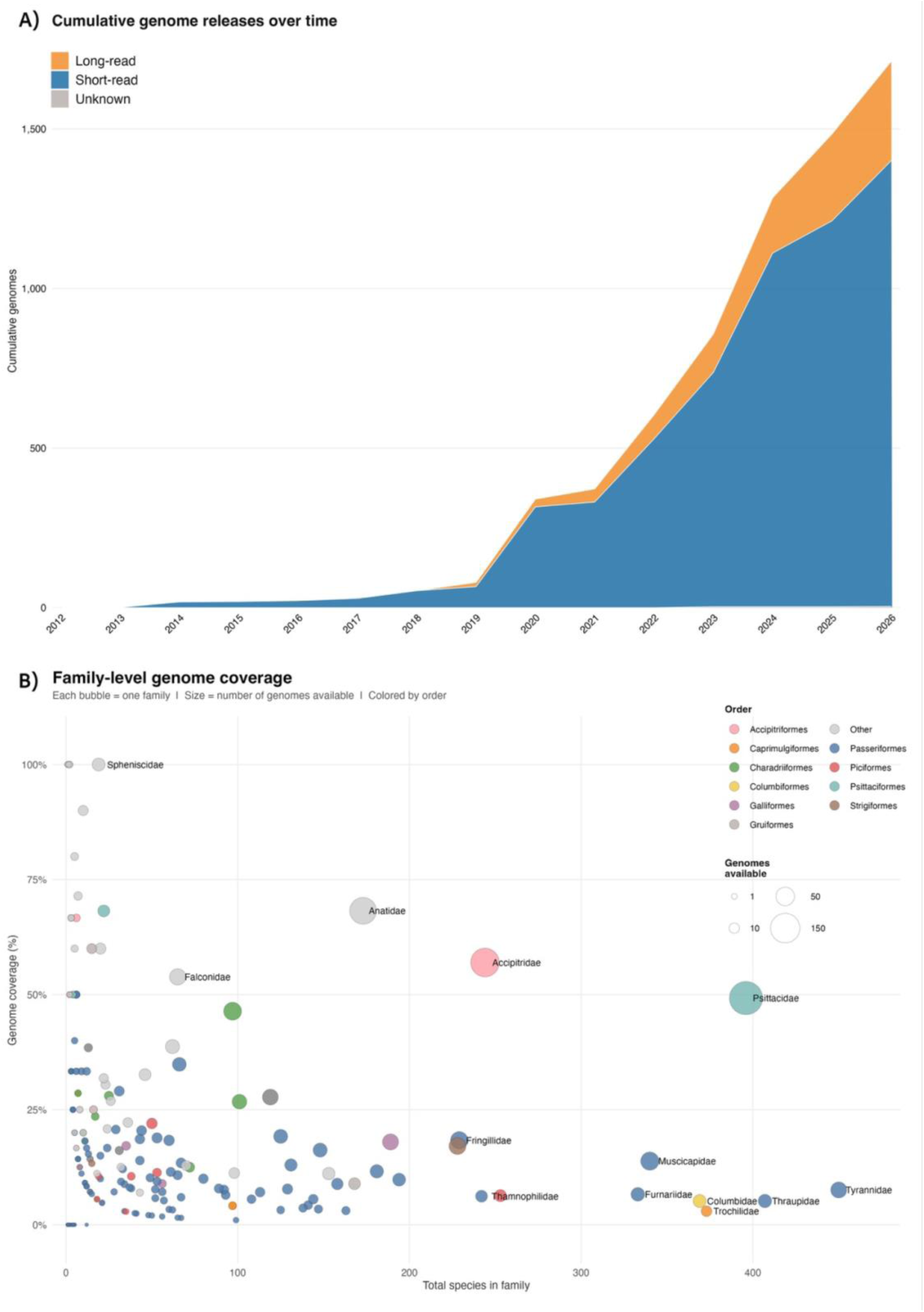
Temporal trends and taxonomic patterns in bird genomes. A) Cumulative bird genome assemblies released over time till March 2026, grouped by sequencing read type. Long-read assemblies (red) contribute an increasing fraction of new genomes in recent years, while short-read assemblies constitute ∼80% of the genome assemblies overall; genomes of unknown read type (grey) make up a negligible fraction of the total (inset, 2020–2026). B) Family-level genome coverage across bird orders. Each bubble represents one avian family, positioned by total species richness (x-axis) and the proportion of species within that family with an available genome assembly (y-axis); bubble size reflects the absolute number of genomes available, and color denotes taxonomic order.

There is a substantial acceleration of genomes produced starting from 2020 (Figure 2A). Pre-2021, the number of genomes across regions was sparse outside temperate regions. In contrast, post-2021 genome assemblies accounted for 80% of all species with genomes, pointing towards rapid sequencing and the influence of large-scale initiatives. This increase led to measurable gains in genomic coverage across parts of South America, Africa, and Southeast Asia. Nevertheless, even after 2021, proportional coverage in most tropical biodiversity hotspots remains low compared to temperate regions.

When comparing taxonomy, each order had at least one genome available (Supplemental Figure S1). Family-level representation was inversely related to species richness (Figure 2B): the most species-rich families were among the most poorly covered proportionally, with the ten largest families — including Tyrannidae (34/450, 7.6%), Thraupidae (21/407, 5.2%), and Trochilidae (11/373, 2.9%) — each represented by genomes for well under 15% of their species. Small families (≤5 species) had a mean proportional coverage of 48.6%, more than three times that of large families (≥100 species; 14.5%). The principal exceptions to this pattern were a small number of large families of longstanding economic, domestication, or research interest, such as Anatidae (118/173, 68.2%), Accipitridae (139/244, 57.0%), and Psittacidae (195/396, 49.2%), which were far better covered than other families of comparable richness.

Nevertheless, 24 families lacked any available genome assemblies, all of which were geographically restricted to the Global South, with distributions concentrated mainly in the Indo-Malay and Afrotropics, and a smaller number in the Neotropics and Caribbean regions. Several of these are species-poor families, such as Pluvianidae, Eupetidae, Hylocitreidae, Hypocoliidae, Pityriasidae, Platylophidae, and Zeledoniidae are monotypic whereas Melanopareiidae, Calyptophilidae, Aegithinidae, and Melampittidae comprise a single genus each. On the other hand, Phaenicophilidae, endemic to the Greater Antilles, comprises 4 genera and 8 species, still lacks any representative genomes (Figure 3). Several of these families not represented by genomes are endemic to a narrow geographic area.

**Figure 3:**
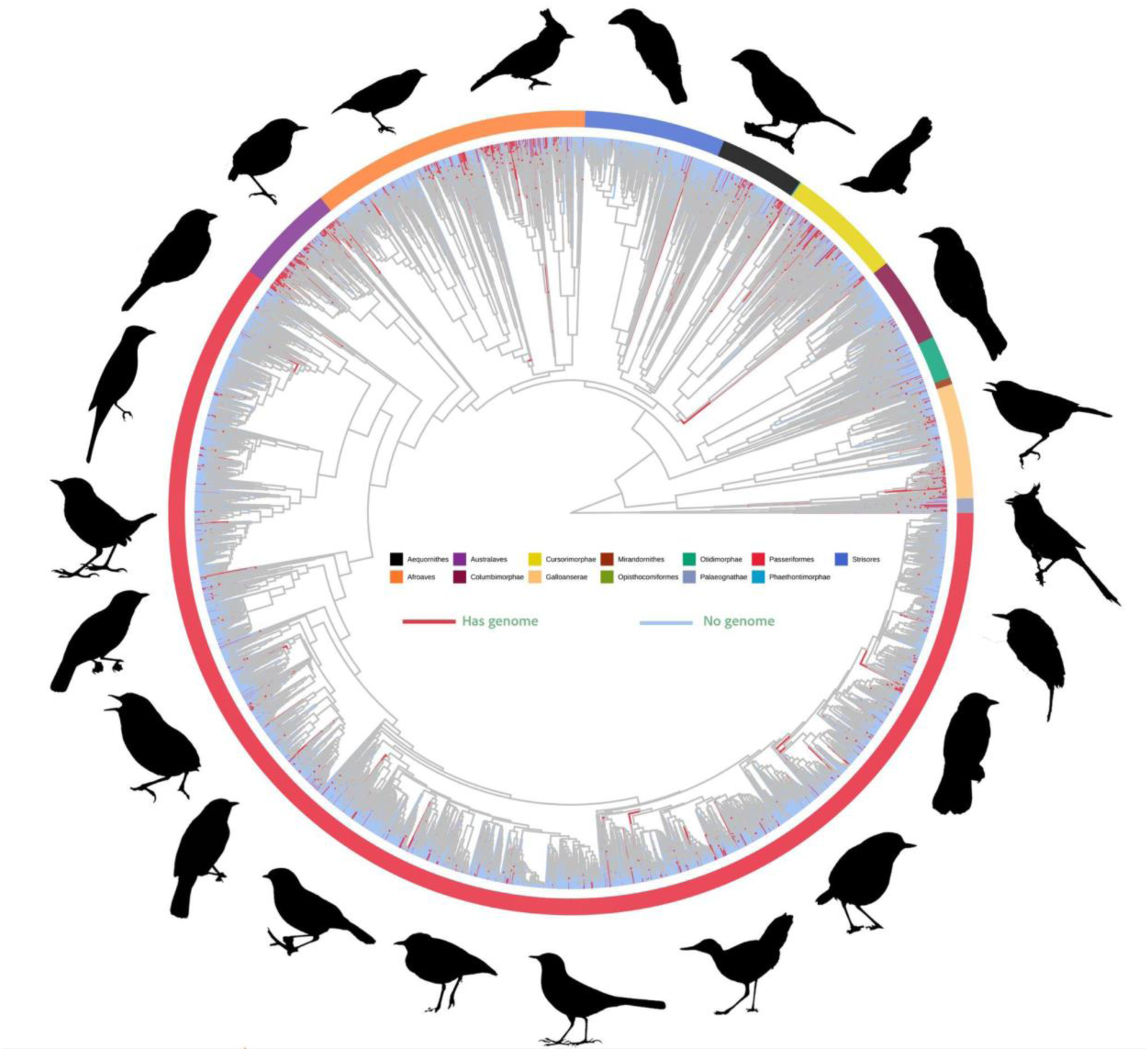
Phylogenetic distribution of avian genomes. Circular phylogram [adapted from McTavish et al., (2025)] of extant bird species, with branches colored to indicate genome sequencing status: red branches represent species with at least one published genome assembly, while light blue branches represent lineages lacking any genome representation. The outer colored ring denotes major avian clades (Aequornithes, Afroaves, Australaves, Columbimorphae, Cursorimorphae, Galloanserae, Mirandornithes, Opisthocomiformes, Otidimorphae, Palaeognathae, Passeriformes, Phaethontimorphae, and Strisores), illustrating how genome sequencing effort is distributed across the avian tree of life. Silhouettes surrounding the phylogeny depict representative bird families with no genome assemblies available, highlighting major taxonomic gaps in genomic coverage despite the overall depth of the underlying phylogeny.

Considering conservation status, genome representation was highest among Least Concern species (1,362/8,742; 15.6%) and declined across categories of increasing threat, with only 38 of 204 Critically Endangered (18.6%), 64 of 395 Endangered (16.2%), and 2 of 165 Extinct species (1.2%) represented by a genome assembly (Figure 4A; Supplemental Figure S2). A comparable gap was evident among EDGE lineages: only 151 of 676 EDGE species (22%) had a genome assembly, leaving 525 (78%) without any genomic resource (Figure 4B). Together, these patterns indicate that threatened and evolutionarily distinctive species, arguably most in need of genomic resources for conservation, are among the least represented in the current dataset.

**Figure 4:**
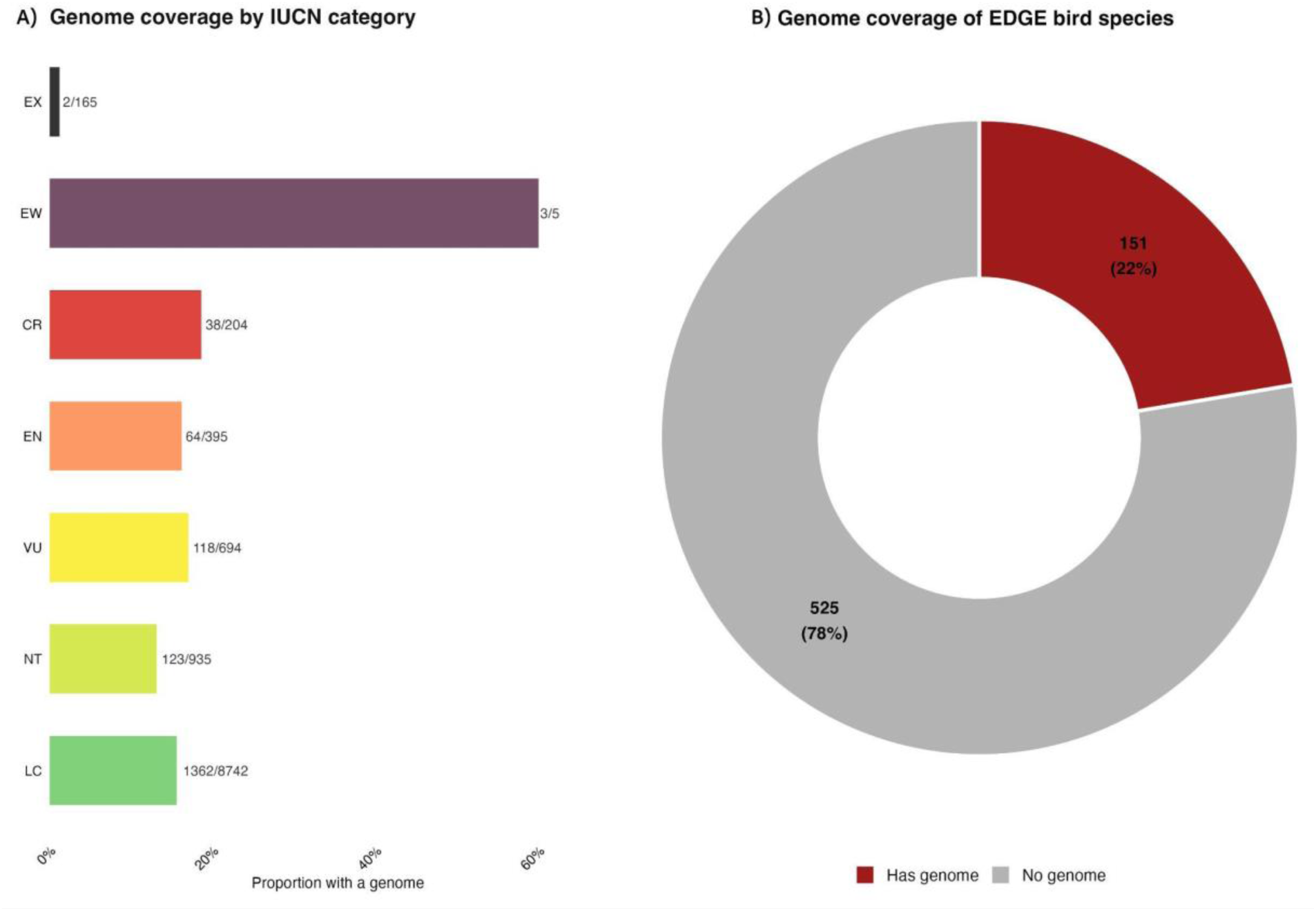
Genome sequencing coverage across avian conservation priorities. A) Proportion of species with at least one genome assembly across IUCN Red List categories (Extinct, Extinct in Wild, Critically Endangered, Endangered, Vulnerable, Near Threatened, Least Concern). Bars represent the fraction of species with genomes within each category, with raw counts annotated as the number of species with genomes / total number of species in each category. B) Genome coverage of EDGE (Evolutionarily Distinct and Globally Endangered) bird species, shown as a donut chart contrasting species with at least one genome assembly (red) versus those lacking any genomic resources (grey). Only 151 EDGE species (∼22%) currently have genome assemblies, while the majority (525 species; ∼78%) remain without genome assemblies.

The global distribution of bird genomes was highly uneven across geographic space (Figure 5). But this unevenness takes a different form depending on whether it is expressed as an absolute count or as a proportion of local richness. In absolute terms, the number of genome-represented species per grid cell was, in fact, highest in the most species-rich tropical regions—particularly the Amazon Basin, which showed the highest per-cell counts in both the pre-2021 and post-2021 periods (Supplemental Figure S3), with the Congo Basin emerging as a secondary hotspot post-2021 (Supplemental Figure S4). This pattern is a direct consequence of underlying variation in species richness: because these regions harbor disproportionately more species overall.

**Figure 5:**
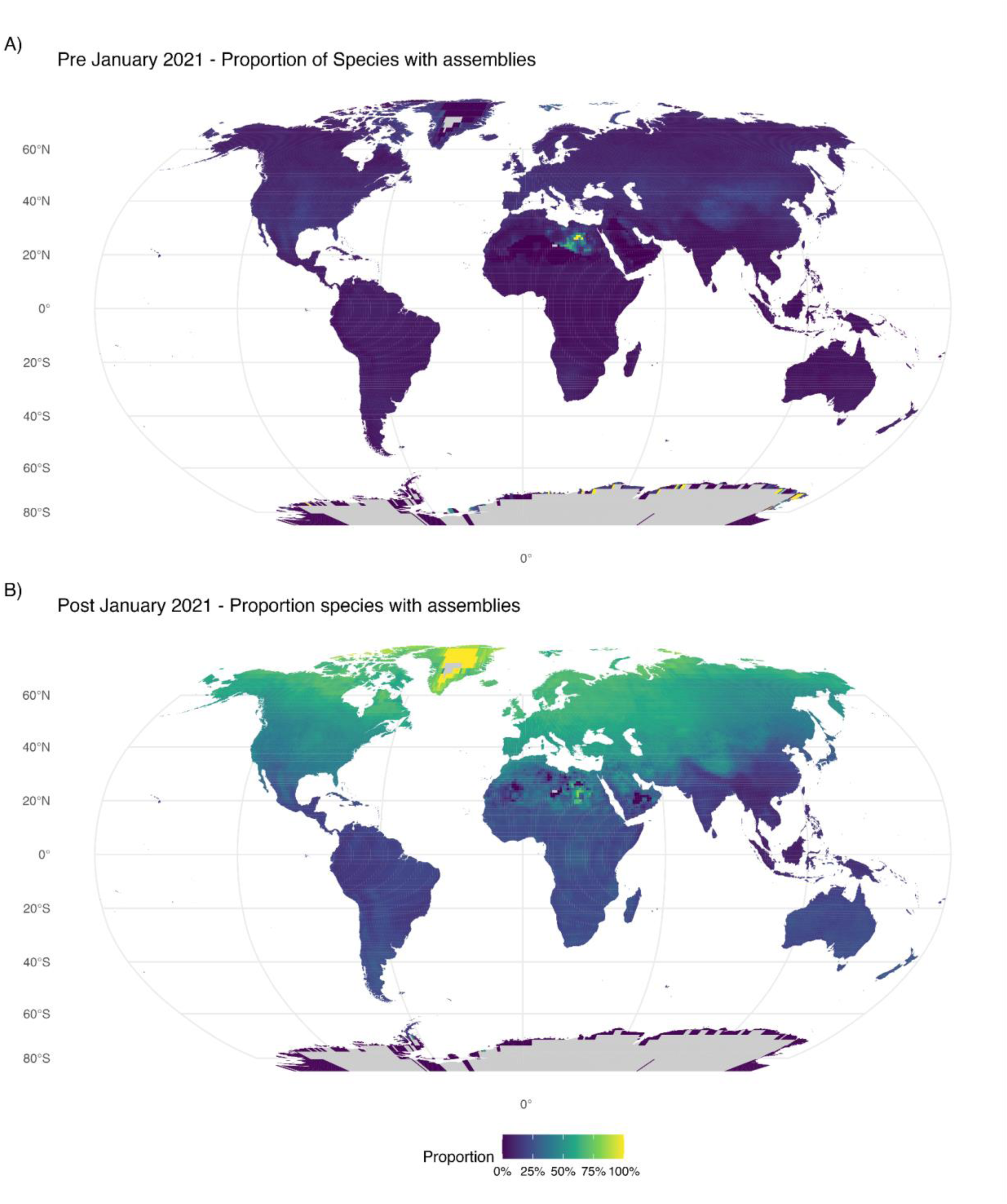
Geographic distribution of avian genomic coverage before and after January 2021. Each cell of a 100 km equal-area grid, clipped to land and displayed in Robinson projection, is shaded by the proportion of bird species occurring in that cell. Species occurrences are breeding and resident ranges (seasonal codes 1 and 2) from the BirdLife/IUCN Bird Species Distribution Maps, and the denominator in every cell is the total number of species whose ranges overlap it. (A) Assemblies released on or before January 2021. (B) Assemblies released after that date, through March 2026. The two panels show non-overlapping release windows rather than cumulative totals, so a species sequenced before 2021 does not contribute to panel B. Grey cells are those with no overlapping species range data.

When expressed as richness-weighted proportional coverage, the disparity becomes unambiguous. Prior to 2021, the proportional representation between GN and GS was relatively low with about two fold difference (Global North: 10.0%; Global South: 4.2%). Following the post-2021 acceleration in genome sequencing (Figure 2A), this gap widened substantially in absolute terms: richness-weighted coverage in the Global North reached 48.0%, while coverage in the Global South rose to only 16.6%. A further contrast emerges from comparing the weighted and unweighted means within each region. In the Global North, weighting by richness raises the mean (48.0% weighted vs 40.3% unweighted), whereas, in the Global South, weighting makes essentially no difference (16.6% vs 16.8%), meaning that its richest cells are covered no better than its most species-poor ones indicating that greater local diversity has not translated into greater sequencing efforts in the tropics.

To test whether the spatial pattern of genomic under-representation in the Global South persists after accounting for conservation status, evolutionary distinctiveness, range and ecology, we modeled genome availability using a generalized linear mixed model with nested taxonomic random effects.

Biogeographic region was a strong and highly significant predictor of genome availability overall (likelihood-ratio test, χ²₅ = 357.3, P < 0.001). After adjusting for conservation status, EDGE species, island endemism, diet, and habitat, species endemic to the Global South had markedly lower odds of genome availability than species endemic to the Global North (OR = 0.23, 95% CI 0.18–0.29, P < 0.001); an approximately fourfold deficit that mirrors the spatial pattern described above. This effect was essentially unchanged when the taxonomic random-effect structure was simplified to order alone (OR = 0.22, 95% CI 0.18–0.27), indicating that the disparity is not driven by taxonomic clustering at the family level. Species belonging to Australasian/Oceanian realms showed a similar odds ratio to that of Global South (OR = 0.21, 95% CI 0.15–0.28, P < 0.001). Trans-regional species, which span both Global North and Global South realms, did not differ significantly from the Global North baseline (OR = 0.93, 95% CI 0.73–1.19, P = 0.59), and Cosmopolitan species, largely globally distributed waterfowl and seabirds, showed substantially higher odds of genome availability (OR = 3.03, 95% CI 1.81–5.07, P < 0.001).

On the probability scale, model-adjusted estimates at reference covariate values corresponded to a 62% (95% CI 49.6–72.9%) chance of genome availability for a Global North species versus 27.6% (95% CI 19.3–37.7%) for Global South (Figure 1b), a gap that persists despite substantial narrowing relative to the raw, unadjusted rates (33.7% vs. 11.2%). The full set of adjusted predictors is reported in Supplementary Figure S5. Conservation status was itself strongly associated with genomic coverage: IUCN-threatened species had lower odds of genome availability than non-threatened species (OR = 0.67, 95% CI 0.52–0.89, P = 0.003), and Extinct/Extinct-in-the-Wild species markedly so (OR = 0.12, 95% CI 0.04–0.32, P < 0.001). The exception was EDGE priority species, which were associated with relatively higher odds of genome availability (OR = 1.38, 95% CI 1.00–1.93, P = 0.051). Island endemism was independently associated with lower genome availability (OR = 0.48, 95% CI 0.39–0.58, P < 0.001). Diet and habitat showed comparatively little association with genome availability once region and conservation status were accounted for, with the exception of a modest increase for the small, heterogeneous “Other” habitat class (OR = 2.62, 95% CI 1.35–5.03, P = 0.004; Supplementary Figure S5, Supplemental Table S1). The full model explained a modest proportion of variance through fixed effects alone (marginal R² = 0.11), but substantially more once taxonomic structure was included (conditional R² = 0.43).

## Discussion

Genome assembly and next-generation data have become central to how we study, compare, and conserve biodiversity, yet the global distribution of these resources remains far from even (Jose, 2026; Linck & Cadena, 2024), and understanding the current gaps and disparities in the genome resources will help in prioritizing future data generation. In birds, this disparity becomes especially important given that they are the best-studied vertebrate group, and they have served as model systems in evolutionary biology and as part of major coordinated large-scale genome initiatives. Despite rapid growth in genome assemblies over the last few years (Figure 2A), current coverage still includes only a small fraction, and substantial gaps remain across geography, taxonomy, and conservation priority. Genomic resources remain strongly uneven and continue to favor the Global North over the Global South. This imbalance persists even after statistically accounting for conservation status, evolutionary distinctiveness, diet, and habitat. Species endemic to the Global South are roughly one-quarter as likely to be represented with a genome than a Global North species, corresponding to an adjusted probability of genome availability of 28% in the Global South versus 62% in the Global North (Figure 1b).

This lack of genome resources matters because genome assemblies are no longer isolated model-organism references. They are essential for phylogenomics, demographic inference, comparative genomics, structural variant discovery, and conservation-oriented population studies. Uneven genomic coverage thus does not simply leave some species unsampled; it shapes the broader biological questions that could be answered and determines which lineages are included in clade-level, genome-wide studies. Our final analysis of currently available data included 1,712 birds with at least one genome assembly, representing only about 15% of the recognized species. This number clearly reflects a recent jump with genome assembly having more than tripled since 2021.

Whole genome representation across all avian orders is an important milestone achieved in 2014 (Jarvis et al., 2014; Zhang et al., 2014), and the field has expanded rapidly, with ∼92% of avian families’ data sequenced by 2020 (Feng et al., 2020). However, this higher-level coverage implies an impression of completeness that upon closer examination shows deep gaps at finer scales – 24 bird families are still lacking genome assemblies, and these absences are not random. Many of these families are monotypic, geographically restricted, species-poor, or phylogenetically distinctive, and all are confined to the Global South. This creates constraints in the comparative analysis of trait evolution, genome structure, and historical biogeography. The gap widens further at the genus level: only 878 of 2,397 recognized genera (∼37.%) currently have a genome assembly. This matters beyond simple taxonomic accounting, because a genome assembly from the same genus provides substantially stronger support for read mapping, variant discovery, and comparative analyses than one from a more distant relative. In this sense, avian genomic resources remain incomplete at every taxonomic scale finer than order, and this incompleteness compounds precisely where it is least visible – among lineages that are both evolutionarily distinctive and concentrated in the Global South.

A similar mismatch can be seen in conservation priorities. Rather than receiving greater genomic attention, proportionally fewer species of conservation concern have been sequenced than non-threatened species – a gradient that tracks conservation urgency in precisely the wrong direction (Figure 4). EDGE species remain largely unsequenced: more than three-quarters (78%) still lack any genomic resource (Figure 4B). Although these species are modestly more likely to have a genome than their threat status alone would predict (Supplemental Figure S5), that signal is nowhere near strong enough to close the gap.

This indicates that current genome assembly efforts are still biased by logistical feasibility, consortium structure, and specimen availability more than by conservation urgency alone. This is perhaps not surprising, since threatened species are often more difficult to sample, more tightly regulated, and less likely to have high-quality tissue samples available. Yet these constraints do not reduce the value or need of genomic resources for conservation; if anything, they strengthen the opportunity for dedicating more effort towards producing these critical resources and more strategically for the species that are most in need.

Much of this pattern is built into the design of the large, coordinated efforts themselves, not a shortfall in how they were executed. Large consortia structured their early phases around phylogenetic representation - first an assembly per order, then per family, and then now per genus, and so on. This is an efficient way to capture maximum phylogenetic and molecular diversity per genome sequenced, and the family-phase of B10K release of 363 genomes spanning over 92% of bird families was a landmark for comparative genomics (Feng et al., 2020). But a scheme optimized for phylogenetic coverage is decoupled from both species richness and geography: it tracks neither where diversity is concentrated nor where species are more threatened, and it tends to undersample rapidly radiating, species-rich clades. We therefore read the near-complete order- and family-level coverage, the persistent genus-level gap (roughly two-thirds of genera), and the weak coupling between genome availability and conservation status as different faces of the same design logic, not as separate problems (Figures 2 and 1b). Consistent with this, genomic coverage tracked accessibility more than either richness or threat: broadly distributed, easily sampled species foremost among them (OR = 3.03, 95% CI 1.81–5.07) showed markedly higher odds of genome availability than range-restricted endemics of the Global South, Australasia/Oceania, or the Antarctic, independent of conservation status. Notably, the consortia themselves have moved in this direction: the EBP now identifies capacity-building in biodiverse developing nations and respect for access-and-benefit-sharing law as preconditions for its own success (Lewin et al., 2022).

Further, the temporal pattern in our results illustrates both the pace of recent progress and the persistence of older inequalities. The marked acceleration of genome assemblies around mid-2020 is consistent with the influence of B10K and other large-scale efforts. This recent expansion, which largely contributed to the first 500 bird genomes by 2021, has clearly improved genomic coverage and increased the number of represented species across several parts of the Global South. At the same time, the broader patterns remain uneven. Proportional coverage across most tropical biodiversity hotspots remains low compared to temperate regions, and the ratio of Global North to Global South coverage widened rather than narrowed over the study period (from 2.4-fold to 2.9-fold; see Spatial distribution of genomes, above) indicating that recent acceleration in sequencing effort has reinforced, rather than corrected, the pre-existing geographic imbalance. This distinction between absolute increase and proportional representation is important because the species-rich tropical and sub-tropical regions may now contain increasing numbers of genome assemblies, yet they remain underrepresented relative to their true species richness. For the Global South the relevant question is therefore not simply whether genomes exist, but whether genomic coverage reflects the true distribution of biodiversity itself.

A further dimension of this imbalance concerns assembly quality rather than assembly count. Contiguous long-read assemblies resolve repetitive regions and structural variation that short-read drafts cannot, but they require high-quality DNA from fresh or cryopreserved tissue (Blom 2021), which has been proven to be difficult to obtain in Tropical species. The field’s shift toward long-read data (Figure 2A) is therefore likely to widen the gap rather than narrow it. The recent direction of the field of genomics towards pangenomes (Fang & Edwards, 2026) further highlights the unevenness in how genomic resources are accumulating. In a few avian taxa, the field has already begun to move beyond single-reference genomes towards population-level resources and pangenomic frameworks, with multiple high-quality assemblies being generated within the same species [For example, Scrub-Jays, Edwards et al., (2025); House Finch, Fang & Edwards, (2024); Barn Swallow, Secomandi et al., (2023), Jeon et al., (2026) and others Rice et al., (2023)]. This is an important and exciting development because it reflects a shift from simply obtaining a reference genome to characterizing genomic variation, yet at the same time it also underscores the broader imbalance. While some species now benefit from increasingly dense genomic data, several avian families still lack even a single genome. This contrast does not diminish the value and importance of the pangenome-scale efforts, rather it highlights how unevenly the benefits of the genomic era persists across bird clades.

Set against this widening within-species/across-species divide, the most encouraging counter-trend is where new sequencing capacity is being built in the first place. Several consortia, projects, and individual groups across Africa (Ebenezer et al., 2022), Asia (*Hong Kong Biodiversity Genomics*, 2024), South America (Vilaça et al., 2024), and other parts of the Global South are helping to improve taxonomic representation. A recent development is that a growing share of this work is led from within the regions of concern. Parallelly, the number of efforts by individual research groups and national institutions working on regionally important taxa is increasing. Recent examples span every major region of the Global South – to name a few, the Philippine Eagle (*Pithecophaga jefferyi*; Hernandez et al., (2026) in Southeast Asia, Przevalski’s partridge (*Alectoris magna*; Wang et al., (2023), multiple *Emberiza* species; Hu et al., (2025) in East Asia, endemic taxa from the Western Ghats of India; (Vinay, et al., 2026a, 2026b), the Southern Ground Hornbill (*Bucorvus leadbeateri*; Patel et al., (2025) and Helmeted Guinea Fowl (*Numida meleagris*; Shen et al., (2021) in Africa. A complementary route is also visible in the Neotropics, where regionally-led teams have generated high-quality assemblies through consortium infrastructure such as the VGP (Cadena et al., 2026; Canesin et al., 2024; Granger-Neto et al., 2025), a model in which the sequencing capacity is centralized, but the scientific leadership and species prioritization are not.

Looking ahead, aligning genomic coverage with avian diversity does not require any single actor to change course dramatically; only that priorities evolve as the field matures. With order- and family-level milestones essentially met, the value of the next bird genome lies less in adding phylogenetic breadth than in filling the gaps such sampling leaves behind, i.e. the remaining genus-level absences, the 24 families without any assembly, and the EDGE species. These gaps do not arise independently of history: current genomic disparities mirrors older patterns of specimen collection, museum curation, research funding, and scientific infrastructure, and closing them will require more than simply pointing existing sequencing capacity at new targets. Weighting future targets by diversity and conservation need, and generating more of this work locally through regional sequencing and bioinformatics capacity (on the model of the AfricaBP Open Institute (*Open Institute*, n.d.) and decentralized EBP nodes (Lewin et al., 2022), and the African-led “3MAG” initiative to sequence three million African genomes over the next decade (Wonkam, 2021), which illustrates how within-region investment can simultaneously build local infrastructure, train a new generation of regional genome scientists, and generate resources at a scale that would build directly on the coverage B10K and allied efforts have achieved.

Generating genomic resources within the Global South, in other words, has value beyond increasing regional representation on a map: it strengthens local technical capacity, reduces dependence on external sequencing, and links genome generation more directly to local conservation priorities. None of this substitutes for equitable authorship, however only about one in six conservation-genetics papers on Global South taxa list a Global South first or senior author (Carneiro et al., 2025; Linck & Cadena, 2024), and given how much of avian genomics still relies on tissue samples held at northern museums, genuine co-design and collaborative use of those collections are what would ensure closing the resource and participation gaps.

## Acknowledgments

VKL thanks Harsha Kumar, Karthik Thrikkadeeri, Oluwaseun Akinsulire, David Vander Pluym, and Eamon Corbett for meaningful discussions during the development of this manuscript.

## Declaration of AI usage

During the analytical process for the manuscript, we used Claude Sonnet 5 and Opus 4.8 to simplify and improve the processing of R scripts. We used Grammarly to improve the readability and clarity, and Paperpile to manage citations, but we did not use Generative AI tools to write this manuscript. We take full responsibility for the text presented here.

## Data Availability Statement

Associated scripts and data for reproducing the analysis are available on GitHub (https://github.com/stachyris/GlobalSouthGenomics).

## Author Contributions

**Vinay K. L.**: Data curation, Formal analysis, Investigation, Methodology, Validation, Visualization, Writing—original draft, Writing—review & editing. **Chiti Arvind**: Conceptualization, Methodology, Writing—original draft, Writing—review & editing. **Naman Goyal**: Conceptualization, Writing—review & editing. **Ashwin Warudkar**: Conceptualization, Writing— review & editing. **Sushma Reddy**: Supervision, Writing—original draft, Writing—review & editing. **V. V. Robin**: Conceptualization, Supervision, Writing—original draft, Writing—review & editing.

