## Supplemental File for "Disparities in avian genomic resources highlight gaps across the Global South"

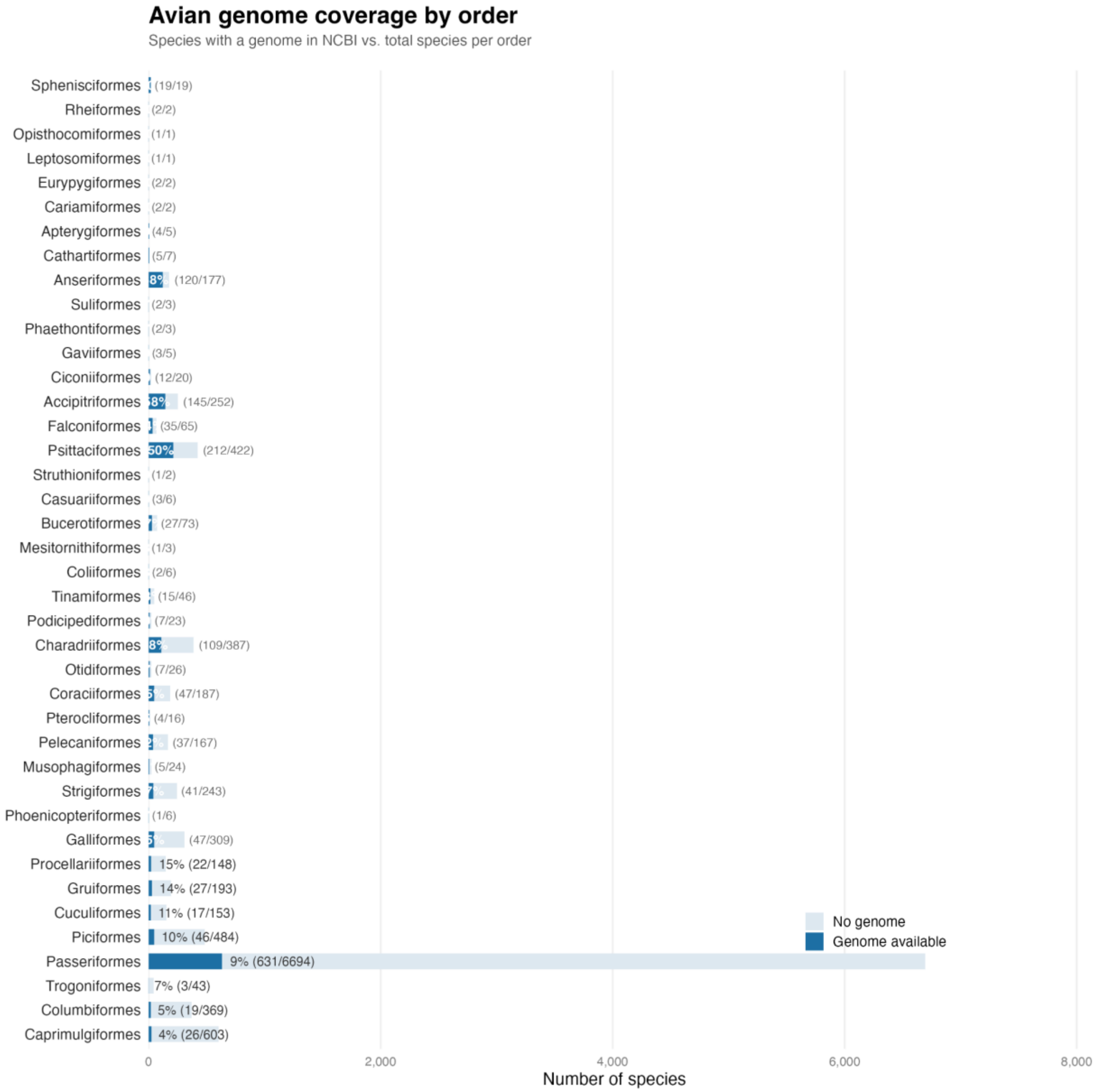


**Figure S1:** Avian genome coverage by order. Bars show species with a genome assembly in NCBI (dark blue) against those without (light blue), ranked by proportion sequenced; labels give percentage and counts (sequenced/total).

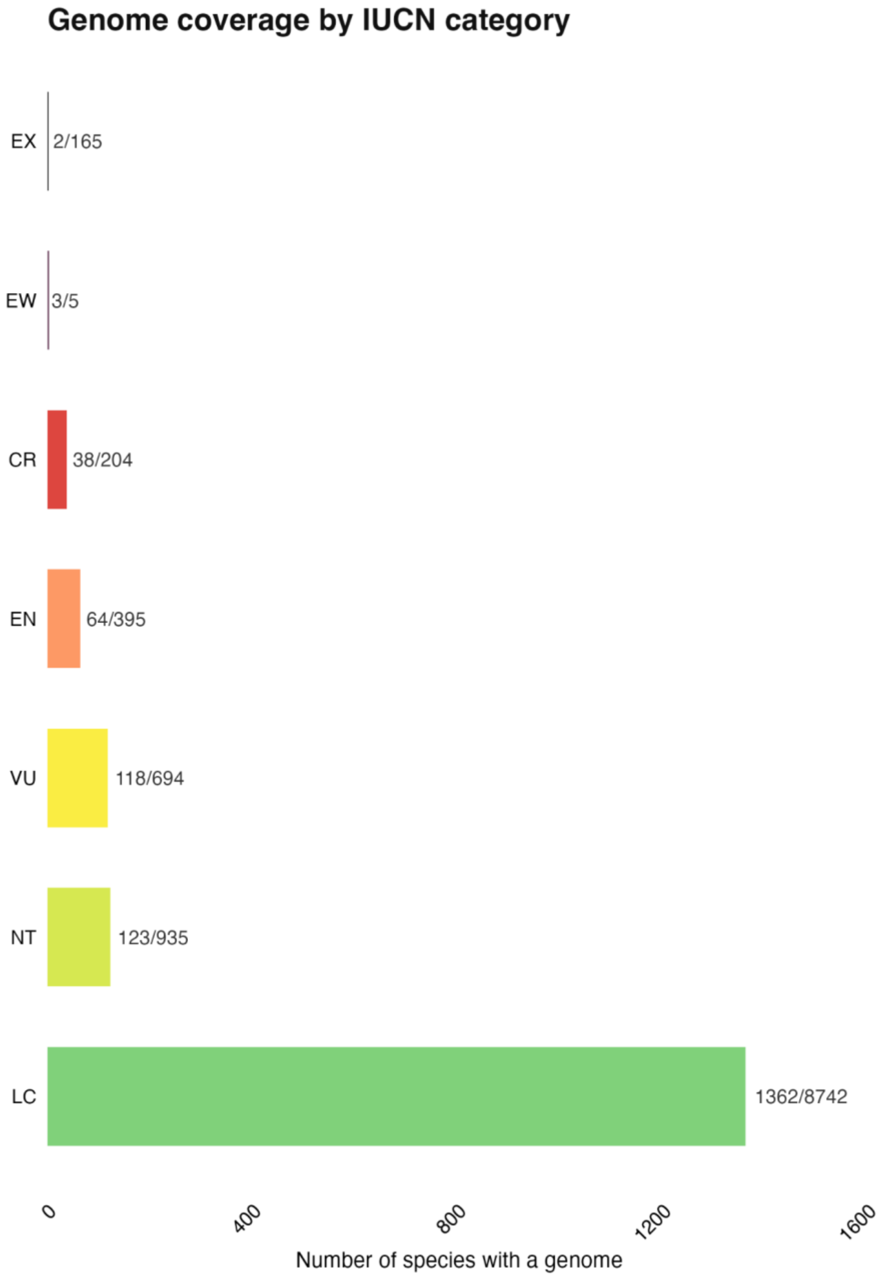


**Figure S2:** Number of species with a genome assembly in each IUCN Red List category; labels give sequenced/total. Bars are absolute counts, not proportions.

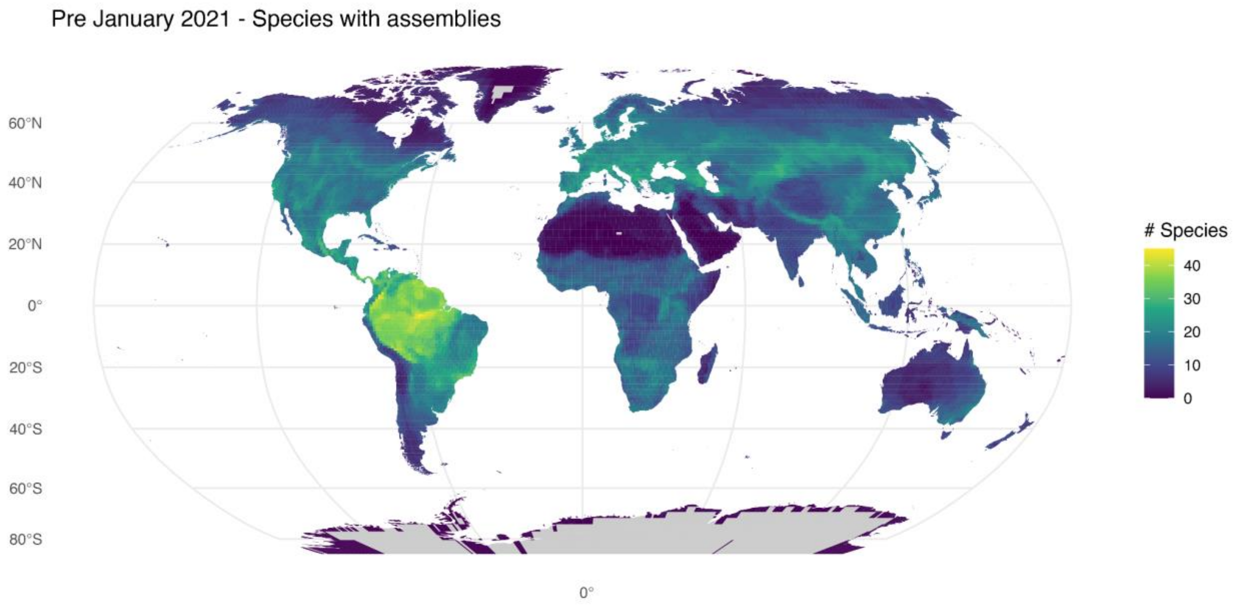


**Figure S3:** Species with genome assemblies released on or before January 2021, per 100 km equal-area grid cell. Ranges are resident and breeding only (seasonal codes 1–2). Grey cells contain no resident or breeding species.

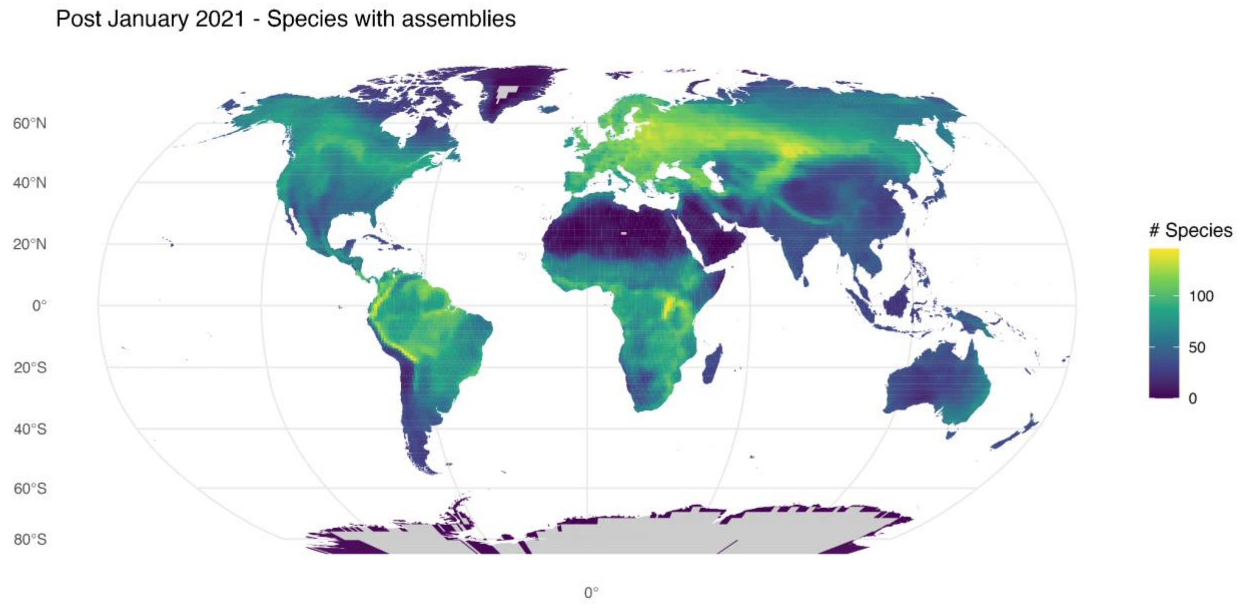


**Figure S4:** Species with genome assemblies released on or after January 2021, per 100 km equal-area grid cell. Ranges are resident and breeding only (seasonal codes 1–2). Grey cells contain no resident or breeding species. Note the colour scale differs from Figure S3.

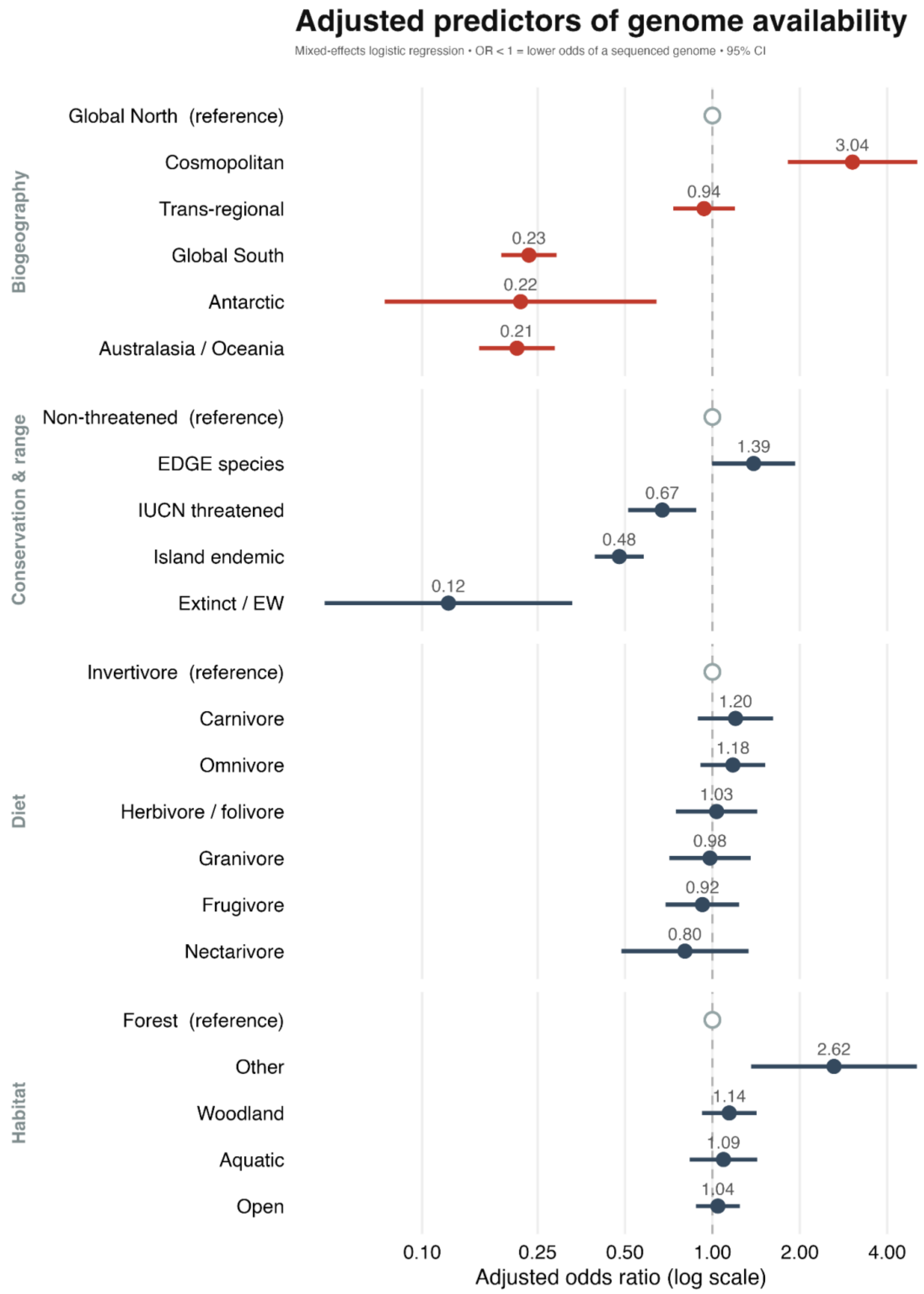


**Figure S5:** Adjusted odds ratios from the mixed-effects logistic regression, with 95% confidence intervals. Open circles mark reference categories (OR = 1): Global North, non-threatened, invertivore, forest. Values below 1 indicate lower odds of a sequenced genome. Estimates are given in Table S1.

**Table S1:** Adjusted odds ratios, 95% confidence intervals, and raw and Benjamini–Hochberg adjusted p-values for the model in Figure S5. Reference categories and random effects are omitted.

| term | OR | lwr | upr | p | p_adj |
|---|---|---|---|---|---|
| regionGlobal South | 0.23 | 0.19 | 0.29 | 1.43E-38 | 2.72E-37 |
| regionAustralasia/Oceania | 0.21 | 0.16 | 0.29 | 5.26E-24 | 5.00E-23 |
| regionTrans-regional | 0.94 | 0.73 | 1.20 | 5.96E-01 | 6.96E-01 |
| regionCosmopolitan | 3.04 | 1.82 | 5.08 | 2.18E-05 | 1.04E-04 |
| regionAntarctic | 0.22 | 0.07 | 0.64 | 5.69E-03 | 1.35E-02 |
| statusThreatened | 0.67 | 0.51 | 0.88 | 3.91E-03 | 1.10E-02 |
| statusExtinct | 0.12 | 0.05 | 0.33 | 2.93E-05 | 1.11E-04 |
| edge | 1.39 | 1.00 | 1.93 | 5.12E-02 | 1.08E-01 |
| isl | 0.48 | 0.39 | 0.58 | 8.30E-14 | 5.25E-13 |
| diet_grpCarnivore | 1.20 | 0.89 | 1.62 | 2.29E-01 | 3.63E-01 |
| diet_grpFrugivore | 0.92 | 0.69 | 1.24 | 5.94E-01 | 6.96E-01 |
| diet_grpGranivore | 0.98 | 0.71 | 1.35 | 9.07E-01 | 9.07E-01 |
| diet_grpHerbivore | 1.03 | 0.75 | 1.43 | 8.39E-01 | 8.86E-01 |
| diet_grpNectarivore | 0.80 | 0.49 | 1.33 | 3.97E-01 | 5.80E-01 |
| diet_grpOmnivore | 1.18 | 0.91 | 1.52 | 2.16E-01 | 3.63E-01 |
| habitat_grpAquatic | 1.09 | 0.84 | 1.43 | 5.19E-01 | 6.96E-01 |
| habitat_grpOpen | 1.04 | 0.88 | 1.24 | 6.23E-01 | 6.96E-01 |
| habitat_grpOther | 2.62 | 1.36 | 5.07 | 4.06E-03 | 1.10E-02 |
| habitat_grpWoodland | 1.14 | 0.92 | 1.42 | 2.26E-01 | 3.63E-01 |
